# Residual Foveal Motion Facilitates Processing of Visually Tracked Objects

**DOI:** 10.1101/2025.09.18.677233

**Authors:** Bin Yang, Jonathan D. Victor, Michele Rucci

## Abstract

Humans and other species visually track moving objects via pursuit eye movements. These movements fail to stabilize the stimulus on the retina, an effect often attributed to errors in oculomotor control. However, it remains unclear whether the resulting residual retinal motion serves visual functions. Using high-resolution eye tracking, we reconstructed foveal motion during discrimination of both stationary and moving stimuli. We show that the retinal motion elicited by pursuit is heavily constrained and strikingly mirrors the motion present during normal active fixation. We then show that this motion performs an important information-compression computational function by equalizing luminance modulations in a low-spatial-frequency range during tracking of natural stimuli. Finally, we show that the resulting visual signals during tracking contribute to perceptual judgments, shifting spatial sensitivity toward lower frequencies relative to fixation. These results suggest that retinal motion during pursuit constitutes an active strategy for encoding space in the joint space-time domain.

## Introduction

The human eyes are never at rest. Even when fixating on a stationary object, the eyes alternate between rapid gaze shifts, known as saccades, and incessant slow movements, known as ocular drifts^1–3^. Yet, we live in a dynamic visual world in which objects are not always stationary but often move. Under these circumstances, humans—like other species—often track moving objects by means of pursuit eye movements, which keep their images within the fovea^4,5^, the retinal region with the highest visual acuity.

Shifting gaze to track moving objects can, in principle, contribute to visual perception through two complementary mechanisms: 1) by regulating visual processing via associated motor—*i.e.*, extra-retinal—signals; and 2) by directly modulating neural activity via the luminance modulations that result from retinal motion. Both possibilities have been extensively studied with stationary scenes. For example, extra-retinal mechanisms associated with saccadic eye movements have been found to suppress visual processing^6–11^ and compress both space^8,12,13^ and time^14,15^. Likewise, an increasing body of literature suggests that, during viewing of stationary scenes, the luminance modulations resulting from eye movements play important functions in visual processing^16–21^.

In the case of pursuit, several studies have focused on the perceptual influences from associated extra-retinal signals. It is established that the perceived motion smear of static background objects is attenuated during pursuit than at fixation, even with identical retinal motion^22,23^, and both the perception of space^24–28^ and time^29^ are distorted during pursuit. More recently, visual sensitivity for chromatic stimuli and achromatic high spatial frequencies has been found to be enhanced by extra-retinal modulations^30–32^.

Despite the extensive research on extra-retinal pursuit influences, however, the potential visual function of retinal motion during pursuit has received little attention. Rather, while it is well-recognized that pursuit eye movements do not fully stabilize the image on the fovea, the remaining motion is commonly attributed to pursuit “errors” that the visual system aims to minimize^33,34^. However, analysis of the foveal motion during pursuit raises the hypothesis that this view may be incomplete. During visual tracking, the eye is often thought to fall behind the target, causing “slippages”, followed by “catch-up” saccades to maintain foveation (Fig. **1**b and d). Crucially, these saccades occur even when the gaze is ahead of the target, raising the possibility that their function is not only to “catch-up”. Thus, the “slippage”-saccade oculomotor cycle resembles the drift-saccade cycle during fixation on steady targets (Fig. **1**a and c) and elicits retinal motion that may effectively stimulate early visual neurons [35–39]. Thus, pursuit “errors” may deliver luminance transients that serve visual functions.

**Fig. 1.**
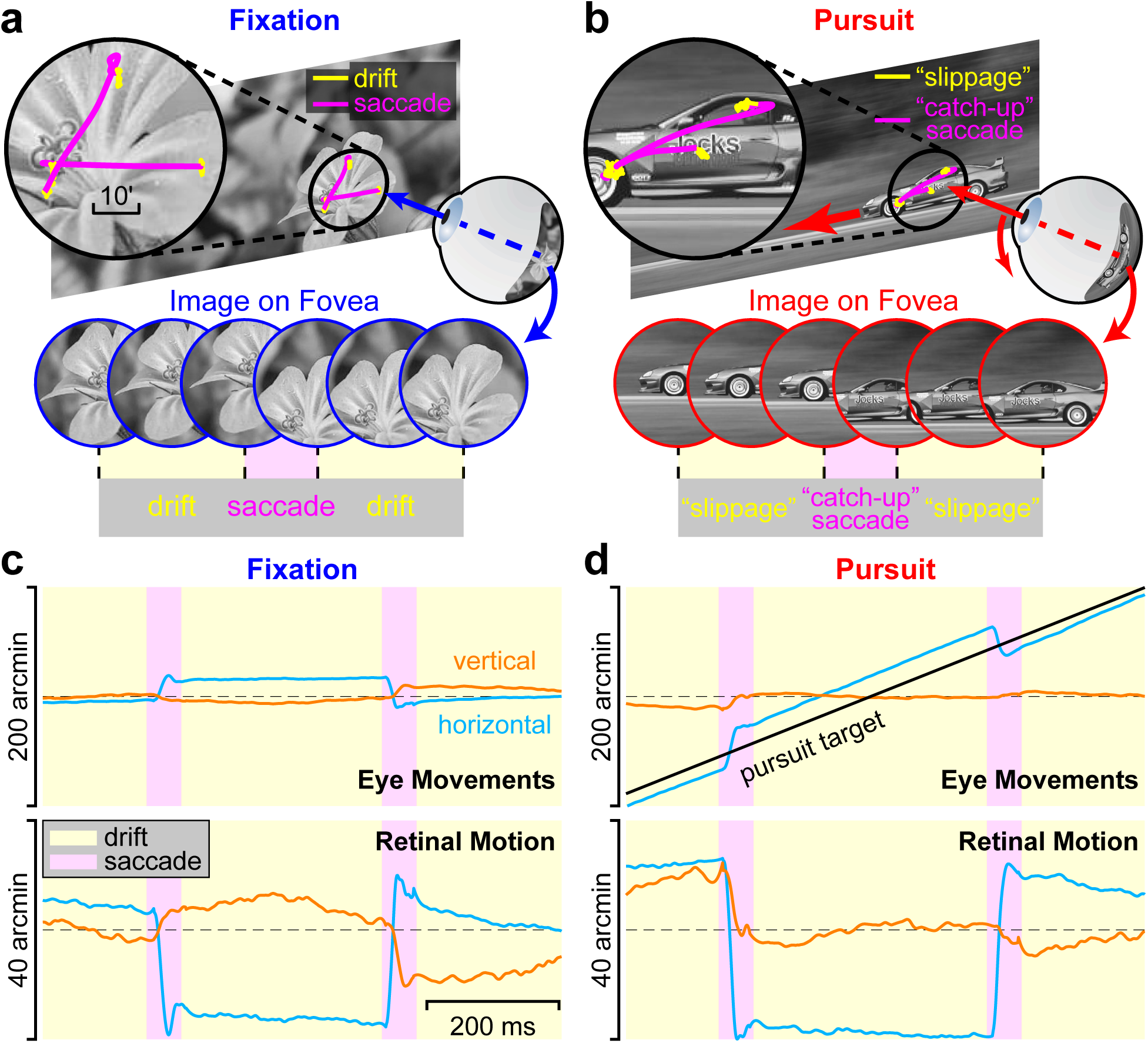
The image on the retina continually moves during both fixation and pursuit. (**a**) During fixation, small saccades (magenta segments) separate periods of smooth motion (ocular drifts; yellow segments) yielding a continually changing stimulus on the retina (sequence of circles). (**b**) The image on the fovea also moves incessantly during pursuit of a moving object, when periods of retinal “slippage” (yellow) alternate with “catch-up” saccades (magenta). (**c**-**d**) Examples of eye traces recorded during fixation and pursuit (top panels) and the resulting motion of the stimulus on the fovea (bottom). Note the similarity in the retinal traces measured in the two conditions (bottom) despite the great differences in eye movements (top).

Here we hypothesize that the pursuit system actively maintains fixation of the target, not to stabilize the target image on the high-resolution fovea, but to leverage retinal motion to facilitate visual processing via luminance transients. To investigate this hypothesis, we compare the retinal motion characteristics of the visual target during both fixation and pursuit. This analysis reveals striking similarities and suggesting shared visual functions. Specifically, pursuit and fixation produce similar patterns of retinal motion across individuals, and idiosyncratic foveation patterns within individuals are preserved across the two oculomotor behaviors. Thus, the two oculomotor behaviors initiate a common mechanism that transforms spatial information into spatiotemporal signals. They thus implement a unified computational principle of active sensation, in which visual processing is facilitated via controlled oculomotor-induced retinal motion.

## Results

### Retinal Image Motion in Pursuit and Fixation

We compared retinal image motion in pursuit and fixation in an orientation discrimination task. Subjects were asked to report whether a Gabor was oriented 45*^◦^* clockwise or counter-clockwise (Fig. **2**a). The Gabor was displayed at the center of a uniform-gray disc slightly lighter than the background, and its contrast slowly ramped up during the course of a trial, to minimize temporal transients from sources other than eye movements. In the “fixation” condition, the stimulus remained stationary at the center of the display. In the “pursuit” condition, the stimulus moved horizontally at a constant velocity of 4 *^◦^/*s across the monitor. Trials under the two conditions were randomly interleaved.

**Fig. 2.**
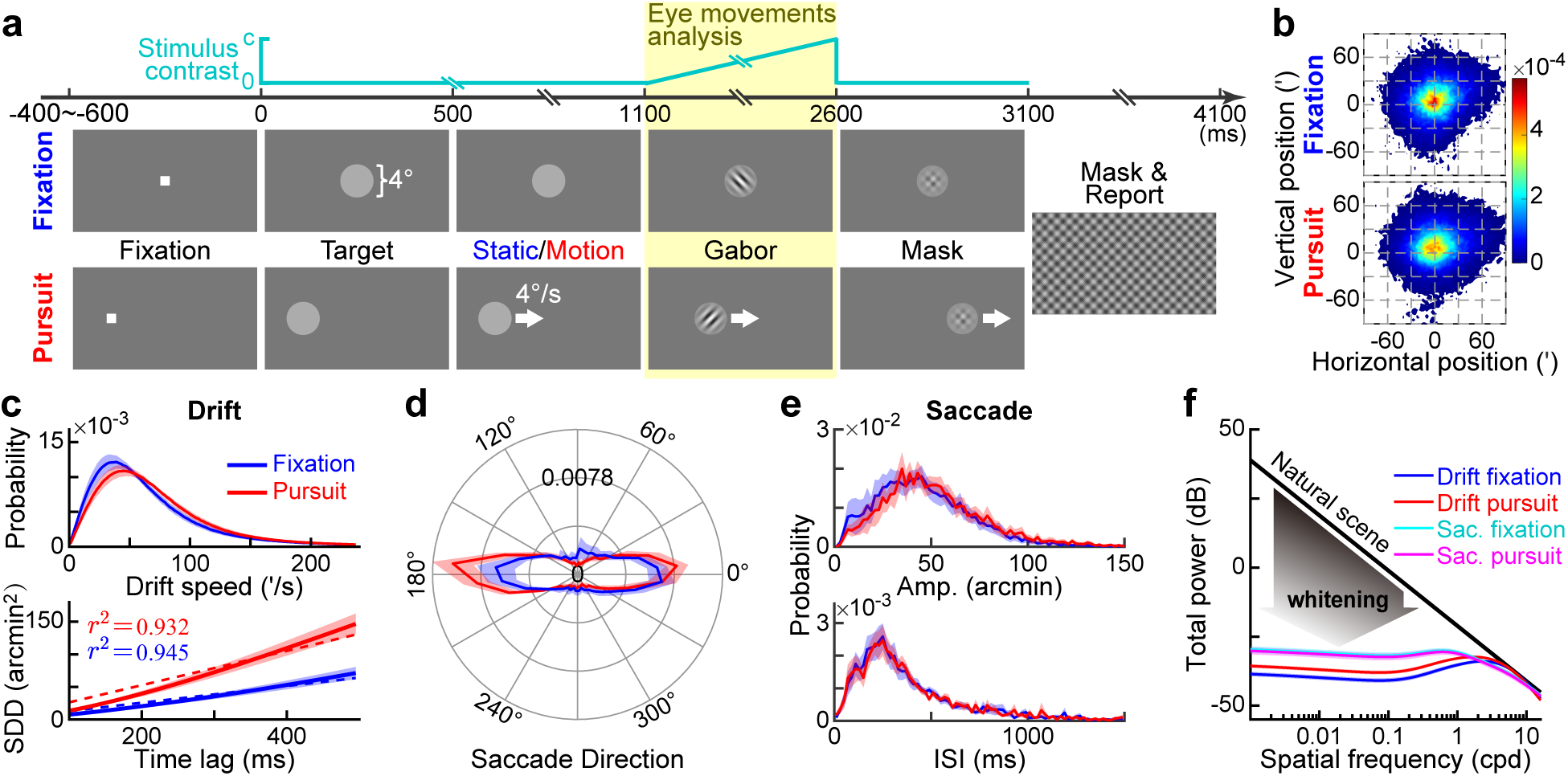
Comparing retinal image motion in fixation and pursuit. (**a**) Experimental paradigm. Subjects were asked to report the orientation (±45*^◦^*) of a Gabor (1 or 10 cycles/degree; *σ* = 0.8*^◦^*) ramping up in contrast (top graph). The stimulus, displayed within a circular aperture, either remained stationary (fixation condition) or moved rightward at 4*^◦^*/s (pursuit). The shaded yellow region indicates the period in which eye movements were compared. (**b**-**e**) Characteristics of eye movements in the two conditions. (***b***) Distributions of gaze positions relative to the center of the stimulus; (***c***) Probability distributions of instantaneous drift speeds (top) and squared drift displacements as a function of time lag (bottom); (***d***) Distributions of saccade directions; (***e***) Distributions of saccade amplitudes (top) and inter-saccadic intervals (bottom). (**f**) In both fixation and pursuit, retinal motion whitens the power spectrum of natural images, yielding luminance signals to the retina with equalized power across a broad range of spatial frequencies. The curves represent the total power delivered by drifts and saccades within the temporal range of human contrast sensitivity^43^ (the curves from saccades in the two conditions overlap). In all panels, data represents averages across *N* =9 subjects. Shaded regions represent ± one SEM.

As expected, subjects maintained steady fixation when the stimulus was stationary and tracked the disc by means of a combination of smooth pursuit and saccades when it moved on the display. On the retina, however, the stimulus was always in motion because of fixational eye movements and imperfect tracking. This motion resulted in similar trajectories of the Gabor stimulus on the retina in the two conditions, both exhibiting periods of slow smooth wandering (retinal drift) interspersed by small saccades. This similarity in retinal motion between the two conditions is evident in the average probability distributions of the retinal displacement of the stimulus: they have the same size and shape, including the way in which they deviate from circularity (Fig. **2**b).

To investigate how different types of eye movements contributed to the similarity in the distributions of Fig. **2**b, we separately compared the retinal motion caused by saccades and drifts. Fig. **2**c shows that retinal drifts were almost identical in the two conditions of fixation and pursuit. In both cases, the instantaneous speed exhibited uni-modal long-tailed distributions (Fig. **2**c, *top*), and the two distributions were closely matched.

Furthermore, the retinal drift during pursuit was well captured by a Brownian motion (BM) model – a model that had previously been shown to apply to fixation^40,41^. To demonstrate this, we examined how the displacement of the stimulus on the retina evolves as a function of time. The bottom panel in Fig. **2**c shows that the squared displacement increased approximately linearly with the time lag in both fixation and pursuit conditions (average *r*^2^ 0.945 and 0.932, respectively; *t*(8) = 1.098, *p* = 0.304, paired *t*-test), indicating that the BM model well describes intersaccadic retinal motion during smooth pursuit. Note that although the BM model fits both behaviors quite well, the effective diffusion constant is higher during pursuit than fixation. We will return to this point below.

The similarity in retinal motion is not restricted to the drifts; it holds for saccades as well. Fig. **2**d and e compare the average characteristics of saccades during fixation and pursuit. The distributions of saccade directions (Fig. **2**d), saccade amplitudes (Fig. **2**e, *top*), and inter-saccadic intervals (Fig. **2**e, *bottom*) were almost identical in the two conditions. Thus, both drifts and saccades contributed to yielding retinal motion with comparable average characteristics during both fixation and pursuit. As we will show below, these characteristics are maintained at an individual level as well, and, moreover, that individuals’ idiosyncratic patterns of gaze and retinal motion are consistent across fixation and pursuit.

Given the results in Fig. **2**b-e, one may expect the retina to be exposed to similar luminance modulations in the two conditions. To investigate the consequence of this motion, we examined how the retinal motion signals measured in our experiments reformat the (1*/f* ^2^ spatial power spectrum^42^) into a spatiotemporal visual flow on the retina. To this end, we estimated how the measured retinal motion redistributes the power of natural scenes across temporal frequencies and computed the resulting strength of visual stimulation within the range of human temporal sensitivity^43^. Previous studies have reported that, with stationary stimuli, both drifts^40,44^ and saccades^45^ equalize (whiten) the power spectrum of natural scenes, yielding luminance modulations with approximately similar amplitudes over a broad range of spatial frequencies. The data in Fig. **2**f show that a similar effect also occurred during pursuit, with drifts whitening up to a higher spatial frequency than saccades. This effect attenuates the statistical redundancies of natural scenes at low spatial frequencies^46,47^ and allows efficient signal transmission through the bottleneck of the optic nerve^47^. Therefore, for both fixation and pursuit, retinal motion transforms visual input signals into a computationally advantageous format. Note, however, that these transformations are qualitatively similar but quantitatively not identical: the reformatting due to retinal drift during pursuit applies a 2-3 dB higher gain, but operates within a somewhat lower frequency range than at fixation. This shift corresponds to the larger effective diffusion constant for drift during during pursuit, than during fixation (Fig. **2**c).

### Idiosyncratic Retinal Motion Patterns across Pursuit and Fixation

As individual differences in fixation behavior are well-recognized^48^, we compared fixation and pursuit behavior at an individual level to determine whether there was evidence that fixational motion and pursuit shared a common oculomotor control mechanism. Specifically, we examined whether patterns of motion were preserved for each individual observer across the two oculomotor tasks. As expected, the amount of retinal image motion varied widely among observers (Fig. **3**a). Strikingly, however, individual subjects exhibited distinctive “fingerprints” of retinal displacement that were preserved across the two conditions. Within each individual, displacement distributions showed a near-perfect match between fixation and pursuit conditions. To quantify the similarity of these distributions between two conditions, and to compare within-subject to between-subject similarity, we used a nonparametric similarity score (1 - the Jensen-Shannon distance; Eq. 1). This metric yielded high within-subject similarity scores (mean ± SD: 0.66 ± 0.06), which far exceeded cross-subject comparisons, whether cross-subject comparisons were between (*t*(79) = 6.806, *p* = 1.8E − 9; two-sample *t*-test) or within each condition (fixation: *t*(43) = 7.205, *p* = 6.5E − 9; pursuit: *t*(43) = 6.597, *p* = 4.9E − 8; two-sample *t*-tests; Fig. **3**b).

**Fig. 3.**
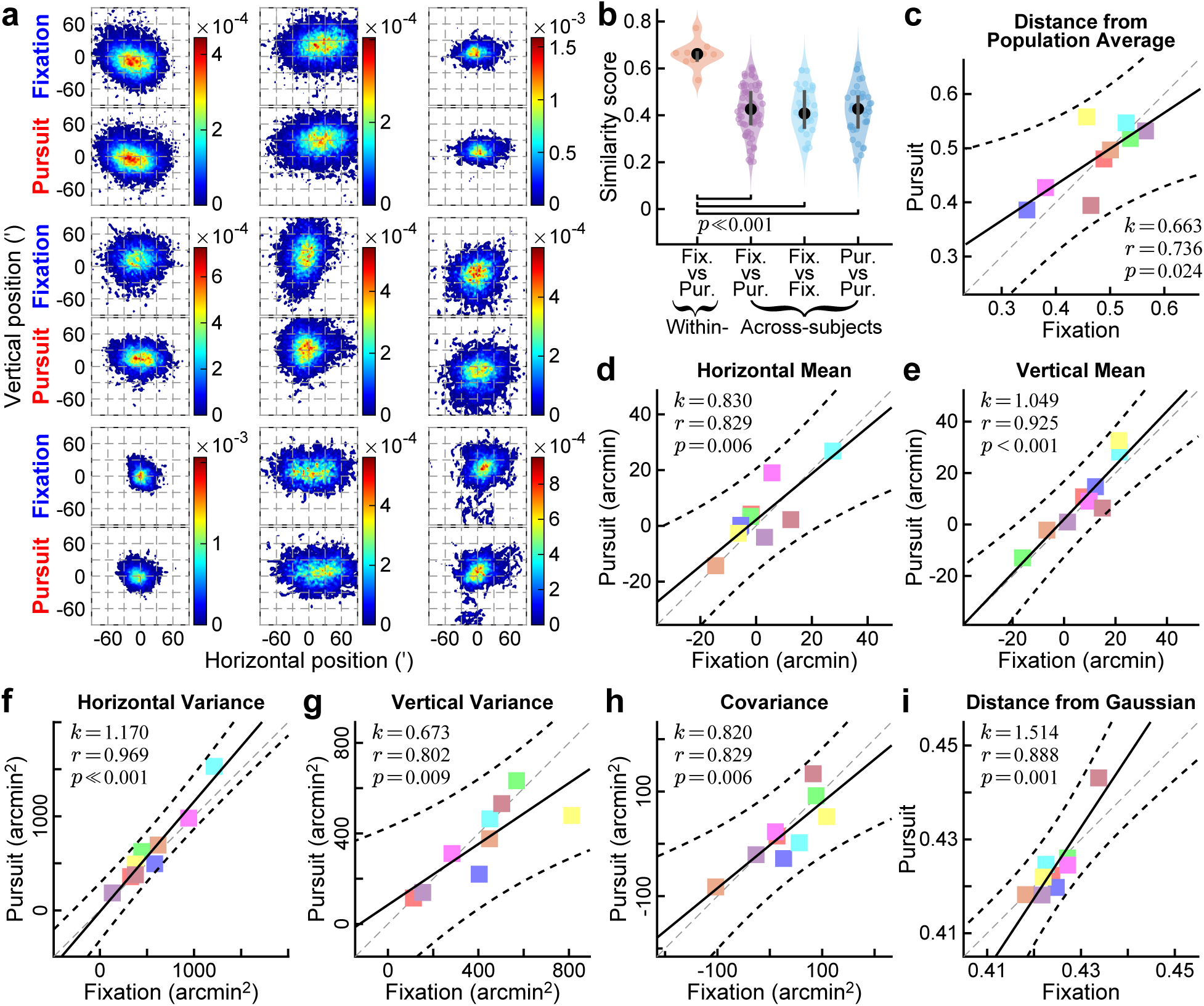
Within-subject consistency of gaze position in fixation and pursuit. (**a**) Gaze probability distributions in fixation and pursuit for each of the *N* = 9 subjects. Data represent offsets in gaze position relative to the center of the stimulus. Note the resemblance of each individual in the two distributions. (**b**) The degree of similarity (Eq. 1) in the fixation-pursuit distributions from the same individual (orange) is significantly higher than that obtained by comparing fixation-fixation (cyan), pursuit-pursuit (blue), or fixation-pursuit maps (pink) from different individuals. *p*-values are the outcomes of two-sample *t*-tests. (**c**) Comparison between how the fixation and pursuit gaze distributions for each individual depart from the population average. The distances (Eq. 2) from the population in fixation and pursuit are strongly correlated (black solid line). (**d**-**h**) Comparing gaze distribution parameters between fixation and pursuit: (*d*) horizontal mean; (*e*) vertical mean; (*f*) horizontal variance; (*g*) vertical variance; (*h*) horizontal-vertical covariance. (**i**) Comparing Jensen-Shannon distances from corresponding 2D-Gaussian distributions. In all panels, *k*, *r*, and *p* denote regression slopes, correlation coefficients, and *p*-values, respectively. Dashed lines represent unity lines (gray) and 95% CIs (black).

Individual deviations from population averages, quantified via Jensen-Shannon distance (Eq. 2), further confirmed this idiosyncrasy resulting in a strong correlation between conditions (*r* = 0.736, *p* = 0.024, linear regression; Fig. **3**c). Moreover, this idiosyncrasy persisted in parametric analyses. We observed robust correlations between fixation and pursuit across all distribution parameters: horizontal and vertical means (Fig. **3**d and e), horizontal and vertical variances (Fig. **3**f and g), and horizontal-vertical covariance (Fig. **3**h). Furthermore, Fig. **3**i reveals a strong correlation in the Jensen-Shannon distance of each empirical gaze distribution from its corresponding 2D-Gaussian fit, defined by that distribution’s own parameters. These results show that each observer maintained their unique way of looking at the stimulus, regardless of whether the stimulus was stationary or in motion.

To investigate the mechanistic basis for these idiosyncratic patterns of retinal image displacement, we compared the individual characteristics of retinal drifts and saccades. Directional distributions of drift segments showed strong within-subject resemblance between fixation and pursuit (Fig. **4**a), with high similarity scores (mean ± SD: 0.93 ± 0.03) far surpassing cross-subject scores (between conditions: *t*(79) = 4.399, *p* = 3.4E−5; fixation: *t*(43) = 4.320, *p* = 9.0E − 5; pursuit: *t*(43) = 3.497, *p* = 0.001; two-sample *t*-tests). In keeping with this result, circular mean directions of drift segments in fixation and pursuit were also strongly correlated across individuals (*r* = 0.828, *p* = 4.8E − 4, rotational regression; Fig. **4**c).

**Fig. 4.**
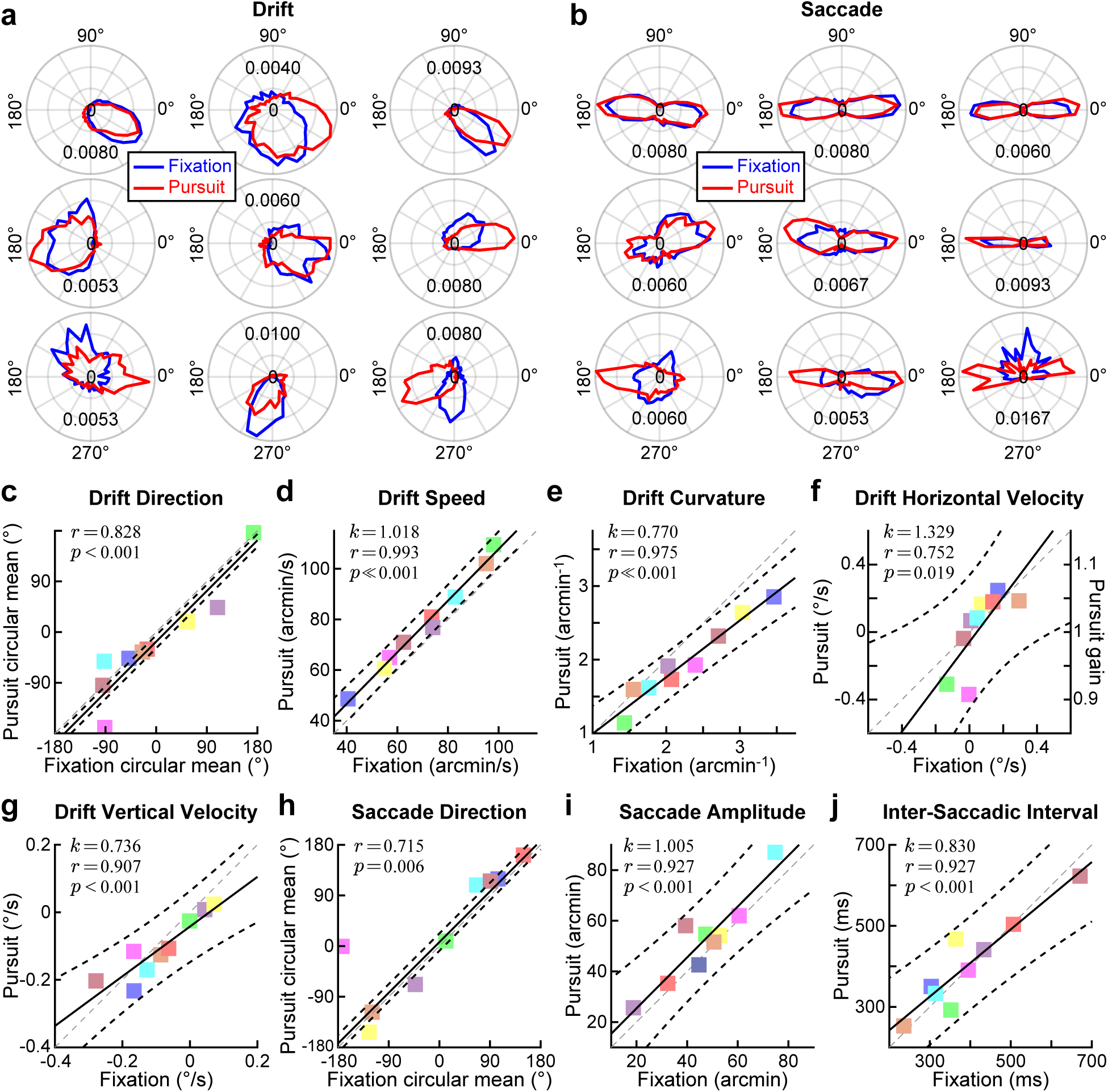
Within-subject consistency of idiosyncracies in retinal motion during fixation and pursuit. (**a**-**b**) Polar probability density distributions of (*a*) drift displacements and (*b*) saccade directions in the two conditions for each participant. (**c**-**g**) Comparison of drift motion: (*c*) Drift direction; (*d*) Instantaneous drift speed; (*e*) Drift curvature; (*f*) Horizontal drift velocity, the right ordinate marks the pursuit gain; (*g*) Vertical drift velocity. Note that while individual motion parameters are highly correlated in the two conditions, retinal drift tends to be faster and less curved in pursuit. (**h**-**j**) Comparison of saccade characteristics: (*h*) Saccade direction; (*i*) Saccade amplitude; (*j*) Inter-saccadic interval. In all *c* − *j* panels, data points represent mean values (circular mean in *c* and *h*) from individual subjects, and graphic conventions are as in Fig. 3c. Note the strong individual correlation of motion characteristics between fixation and pursuit.

For all observers, drift speeds differed somewhat between conditions, an effect that we examine in depth in the following section. Yet, strong individual correlations between conditions were present in both the mean instantaneous drift speed (*r* = 0.993, *p* = 1.0E − 7; Fig. **4**f) and average median curvature of drift segments (0.975, *p* = 8.4E − 6; Fig. **4**g). In fact, both horizontal (Fig. **4**d) and vertical components of drift velocity (Fig. **4**e) showed robust correlations between the two conditions (*r* = 0.752 and 0.907, *p* = 0.019 and *p* = 7.3E − 4; linear regressions). Notably, the horizontal velocity resulted in individual pursuit gains that were close to unity (mean ± SD: 1.006 ± 0.056; *p* = 0.762, *t*-test against unity). Considering the relation between horizontal velocity and pursuit gain (left and right ordinates in Fig. **4**d), these results also imply that each subject’s pursuit accuracy could be predicted from his/her oculomotor behavior at fixation.

Saccadic patterns reinforced this consistency of individual idiosyncracies across fixation and pursuit. The directional distributions of saccades (Fig. **4**b), showed prominent within-subject resemblance between fixation and pursuit (mean ± SD of similarity scores: 0.942 ± 0.033), significantly exceeding the comparisons across subjects (fixation-pursuit: *t*(79) = 4.065, *p* = 1.1E−4; fixation: *t*(43) = 4.023, *p* = 2.3E−4; pursuit: *t*(43) = 4.161, *p* = 1.5E−4; two-sample *t*-tests). Interestingly, catch-up saccades^49^ were not predominant in our experiments, with saccades during pursuit directed both towards and against the direction of the moving target. Moreover, circular mean directions (Fig. **4**h), saccade amplitudes (Fig. **4**i), and inter-saccade intervals (Fig. **4**j) all exhibited strong correlations between fixation and pursuit (circular mean direction: *r* = 0.715, *p* = 0.006, rotational regression; for both saccade amplitudes and inter-saccade intervals: *r* = 0.927 and *p* = 3.2E − 4, linear regressions). These results show that the way saccades affected retinal image motion during pursuit was very similar to that of fixational saccades.

Overall, retinal image motion during pursuit exhibited striking similarities to that observed during fixation, both in population-level analyses and individual subjects. This striking resemblance suggests that retinal motion patterns represent an intrinsic, subject-specific feature that remains consistent across oculomotor tasks. Such findings challenge traditional perspectives that interpret retinal motion during pursuit as mere artifacts of pursuit inaccuracies.

### Retinal Image Motion Accounts for Visual Performance

While participants maintained qualitatively similar retinal motion patterns across conditions, systematic quantitative differences emerged in key retinal drift characteristics. Despite the strong fixation-pursuit correlations in drift speed and curvature shown in the previous figures (Fig. **4**f and g), both parameters differed between the two conditions. The mean instantaneous drift speed was significantly higher during pursuit (mean ± SD: 70.9 ± 19.1 arcmin/s) than during fixation (78.2 ± 19.6 arcmin/s; *t*(8) = 9.143, *p* = 1.7E− 5, paired *t*-test; Fig. **5**a), whereas the median drift curvature was significantly smaller during pursuit (*t*(8) = 4.498, *p* = 0.002, paired *t*-test; Fig. **5**b). These differences resulted in a more rapid and diffuse motion during pursuit, which is quantified by doubling the diffusion constant, a parameter that describes how much the eye moves in a Brownian motion^40,41^ model of ocular drift (mean ± SD: 32.2 ± 11.7 vs 66.0 ± 20.6 arcmin^2^/s; *t*(8) = 5.635, *p* = 4.9E − 4, paired *t*-test; Fig. **5**c).

**Fig. 5.**
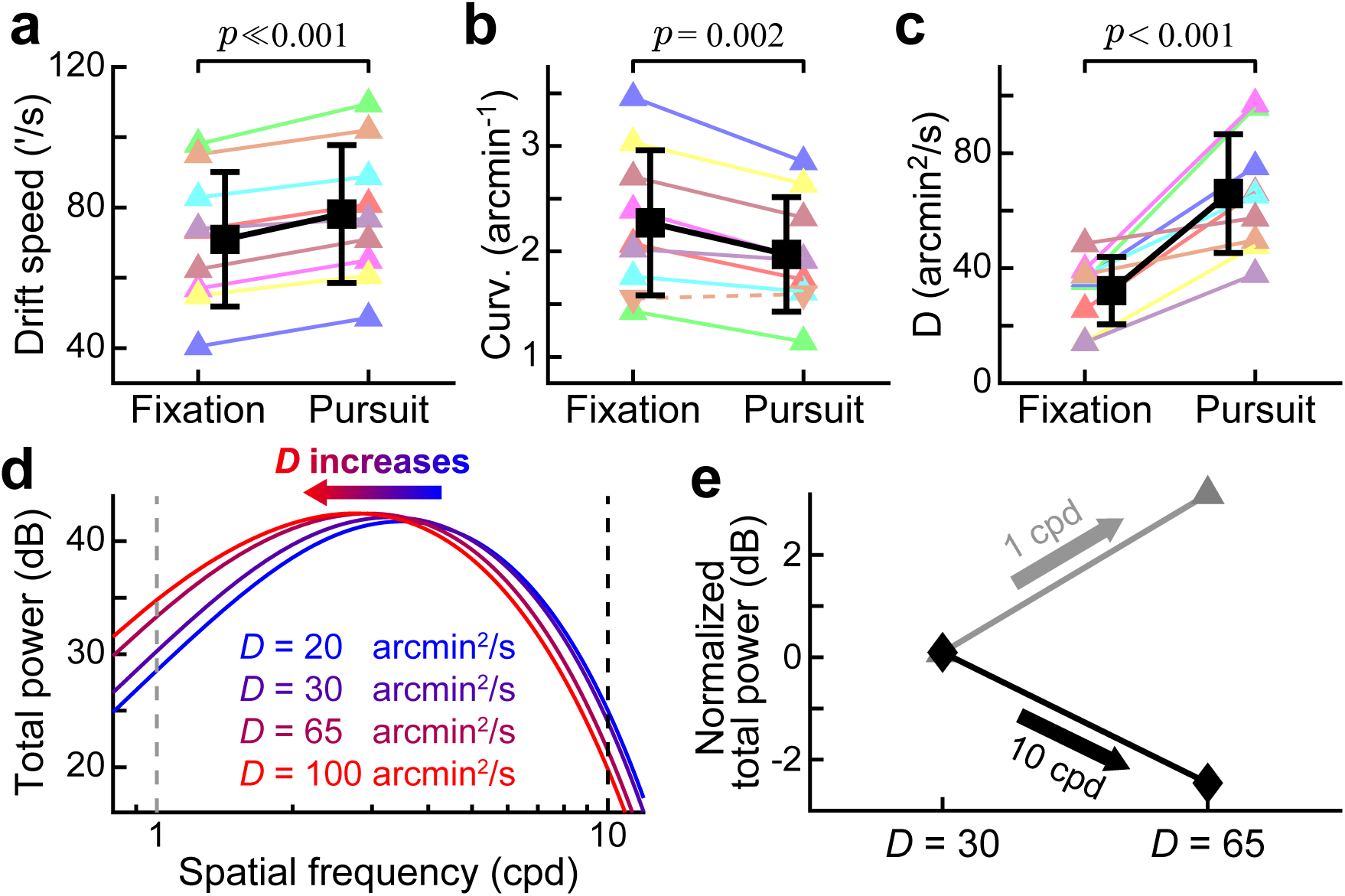
Differences in retinal drift characteristics and their expected perceptual consequences. (**a**-**c**) Direct comparison of retinal drift motion in fixation and pursuit: (***a***) mean instantaneous speed; (***b***) mean instantaneous curvature; (***c***) corresponding diffusion constant. In each panel, both averages across subjects (black symbols) and individual data (colored symbols) are shown. *p*-values are results of paired *t*-tests. All individuals comparisons are statistically significant (*p* ≤ 0.05, one-tailed Wilcoxon rank-sum tests), except for one subject’s curvature data (downward triangle and dashed line in ***b***). (**d**) Within the range of human temporal sensitivity^43^, the luminance modulations delivered by a more diffuse drift shifts power toward low spatial frequencies (arrow). (**e**) With the diffusion constants corresponding to the trajectories measured in our experiments, this effect should result in a stronger visual input signal at 10 cpd and a weaker input at 1 cpd (dashed lines in ***d***).

This quantitative difference provides an opportunity to examine the visual functions of retinal drift during pursuit, as it enables quantitative predictions of the consequences of oculomotor-induced temporal modulations on the retina. Fig. **5**d and e show how increasing the diffusion constant, *D*, in a Brownian motion model of ocular drift affects the power of these induced luminance modulations. The curves in Fig. **5**d represent the total power within the temporal sensitivity range of the human visual system^43^ parametric in *D*. As shown by these data, the strength of visual input signals is not uniform across spatial frequencies, but follows a band-pass function that peaks at a frequency determined by *D*. Increasing the diffusion constant shifts the band of strongest stimulation towards lower spatial frequencies, thereby enhancing low-frequency input signals at the expenses of high spatial frequencies (Fig. **5**e). This effect is expected to shift sensitivity from high to low spatial frequencies in the presence of the larger motion observed during pursuit.

To test these predictions, we estimated the strength of the visual input signals experienced by the subjects in the two experimental conditions. This was achieved by reconstructing, for each participant, the visual input caused by the retinal drifts measured in the experiment and estimating the resulting power within the range of human temporal sensitivity. Consistent with the Brownian motion approximation (Fig. **5**e), changes in the retinal drifts during pursuit and fixation influenced the strength of visual input signals. During pursuit, subjects experienced significantly stronger input signals with the low-spatial-frequency stimulus (*t*(8) = 4.492, *p* = 0.002; paired *t*-test) and weaker input signals with the high-frequency stimulus (*t*(8) = 2.625, *p* = 0.030, paired *t*-test; Fig. **6**a). Individually, all subjects exhibited stronger signals during pursuit of the low-spatial-frequency stimulus, a difference that was statistically significant in all but one participant (*p <* 0.001, one-tailed Wilcoxon rank-sum test). Similarly, lower power during pursuit of the high-spatial-frequency stimulus was observed and statistically significant in all but one subject (*p <* 0.022, one-tailed Wilcoxon rank-sum test).

**Fig. 6.**
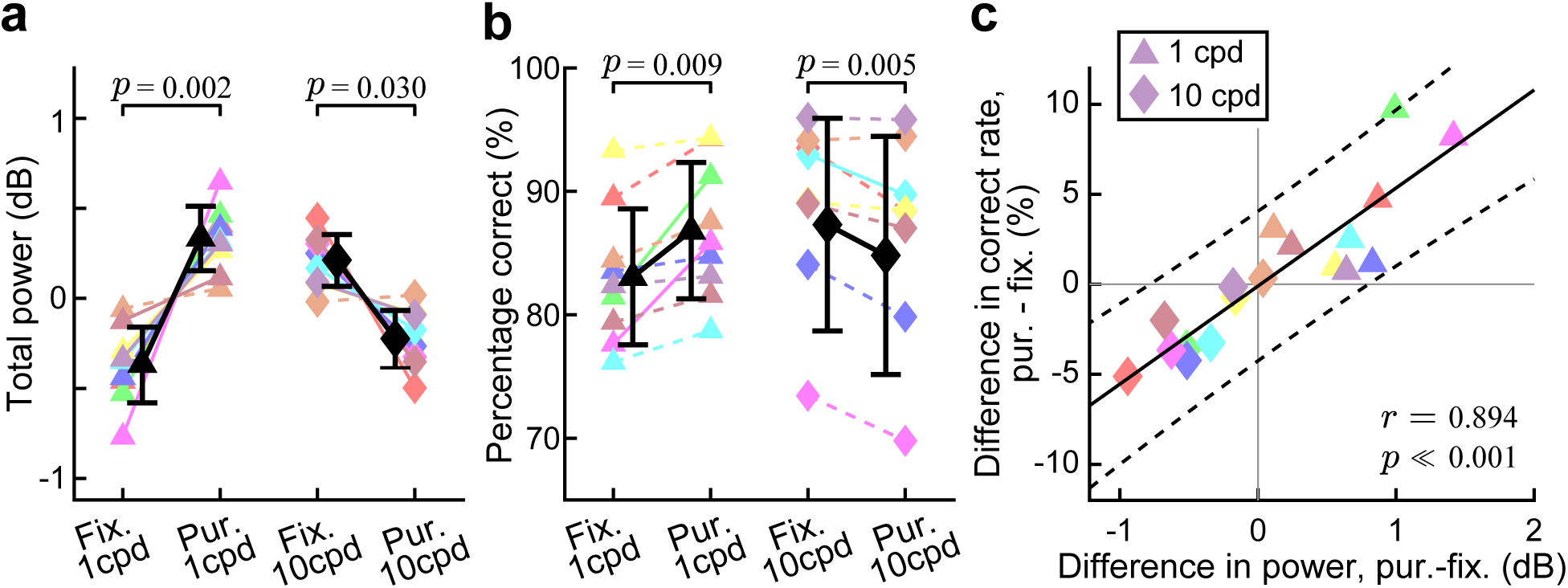
Luminance modulations delivered by retinal drift predict perceptual performance. (**a**) Power delivered by retinal drift within the temporal range of visual sensitivity during fixation and pursuit. Colored symbols show data from individual subjects, each normalized by their overall mean across conditions. Solid lines mark significant differences (*p <* 0.05, one-tailed Wilcoxon rank-sum test). Black symbols represent group averages ± one SD. Power was higher at 1 cpd during pursuit and at 10 cpd during fixation (paired *t*-tests). (**b**) Comparison of performance in the two conditions. Graphic conventions are as in *a*. (**c**) Correlation between the strength of visual input signals and performance. Symbols mark data from individual subjects (different colors) at both 1 cpd (triangles) and 10 cpd (diamonds). Solid and dashed black lines represent the linear regression and its 95% CIs.

Visual performance closely followed the strength of the input signals delivered by retinal drift. Performance during pursuit improved with the 1-cpd stimulus and worsened with the 10-cpd stimulus (Fig. **6**b). These differences were highly consistent across subjects, resulting in strongly significant effects (*t*(8) = 3.450 and 3.851, *p* = 0.009 and 0.005, for the low and high spatial frequencies, respectively; paired *t*-test). All 9 subjects exhibited enhanced performance when tracking the low spatial frequency stimulus, while 8 out of 9 demonstrated impaired performance when tracking the high spatial frequency stimulus. Moreover, individual perceptual differences between the two conditions were strongly correlated with the changes in power experienced by each observer (*r* = 0.894, *p* = 5.7E − 7; Fig. **6**c), so that the subjects who experienced the strongest differences in the strengths of luminance modulations were also those who exhibited the most marked perceptual influences. Thus, perceptual modulations were well predicted by the retinal input signals experienced by each individual observer.

Having found that retinal drift effectively modulates both retinal input strength and visual sensitivity, we also examined possible influences from saccades. Our earlier analyses (Fig. **2**e and Fig. **4**i and j) show that saccades maintain similar characteristics—both amplitudes and rates—in fixation and pursuit, therefore likely contributing in similar ways in the two experimental conditions. In both conditions, however, the number of saccades varies across trials, with some trials containing more saccades than others. This variability implies that saccades may differentially influence trials depending on their frequency.

Fig. **7**a and b illustrate how saccades influence the strength of visual input signals. In this example, saccades are modeled as sigmoidal functions of time (Eq. 8), which interrupted an otherwise continuous Brownian drift motion. Consistent with previous reports^45,50^, because of their larger displacements relative to drifts, saccades primarily affect low spatial frequencies (Fig. **7**a). Thus, when observing a 1-cpd stimulus, the occurrence of a saccade increases the power delivered within the temporal range of visual sensitivity. In contrast, for a 10-cpd stimulus, the strength of visual signals remains largely unchanged regardless of the presence or absence of saccades (Fig. **7**b).

**Fig. 7.**
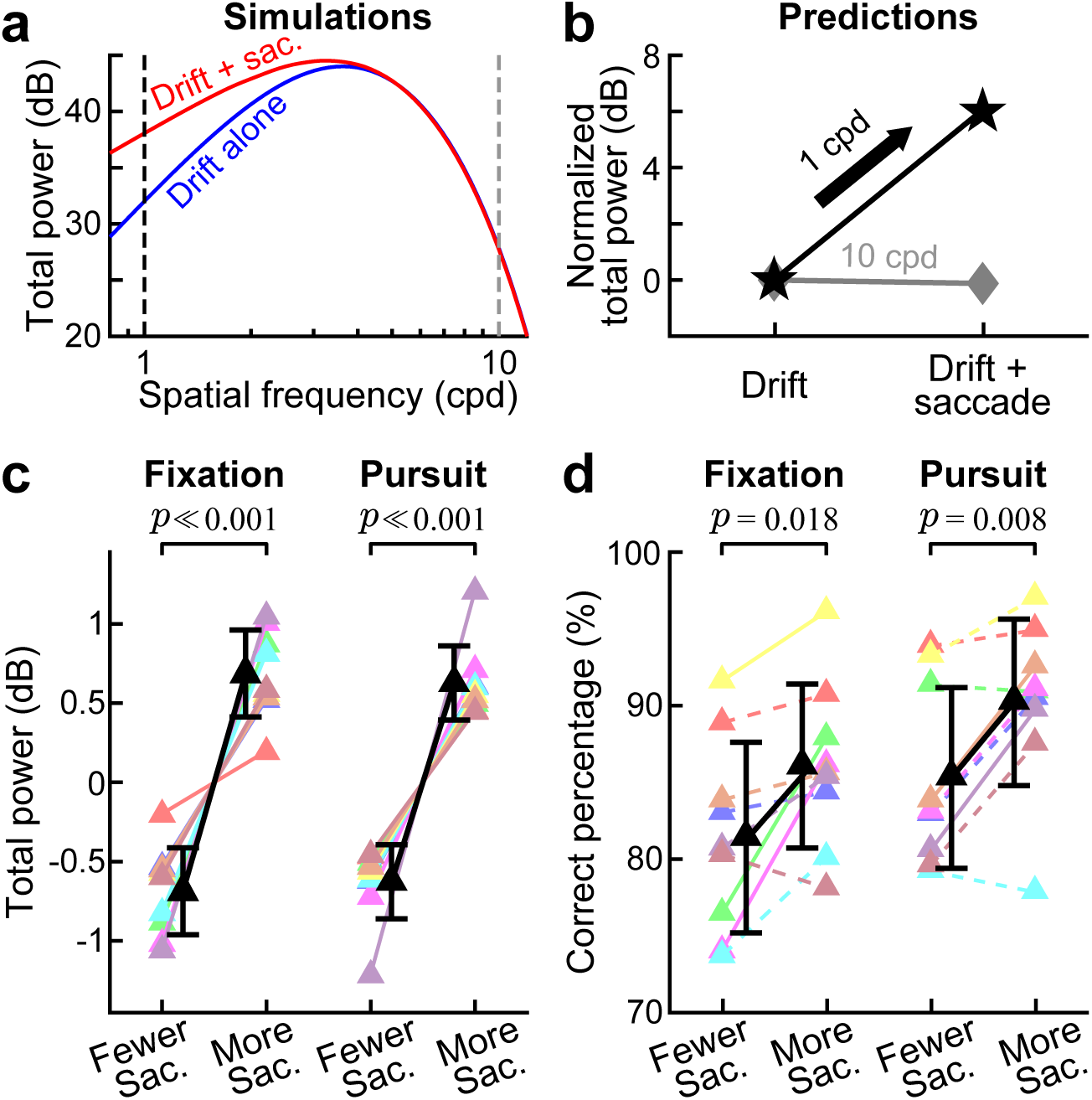
Perceptual contributions of saccades. (**a**) Simulated retinal input power within the human temporal sensitivity range^43^, resulting from drift alone (modeled as Brownian motion) and from drift interrupted by a saccade (simulated with a sigmoid function). (**b**) These simulations predict that saccade occurrence enhances the retinal input signal at 1 cpd, with little effect at 10 cpd (dashed lines in ***a***). (**c** and **d**) Experimental data at 1 cpd reveal differences in retinal stimulation strength (*c*) and perceptual performance (*d*) between trials with above- and below-median saccade counts. Graphic conventions as in Fig. 6.

This low-frequency enhancement was also present in the reconstructions of the visual input signals based on the measured saccades of the participants (Fig. **7**c). Given the similarity in their characteristics, saccades delivered comparable power during fixation and pursuit (*t*(8) = 1.209, *p* = 0.261). However, in both conditions, trials with above-median saccade counts possessed significantly enhanced retinal signals at low frequencies (fixation: *t*(8) = 7.506, *p* = 6.9E − 5; pursuit: *t*(8) = 8.029, *p* = 4.3E − 5; paired *t*-tests), an effect significant in all nine participants (*p* ≤ 8.6E − 5, one-tailed Wilcoxon rank-sum tests).

Mirroring this enhancement in visual input, reporting accuracy for the l cycle/degree stimulus also increased with saccade frequency (Fig. **7**d). In both fixation and pursuit, subjects were more likely to correctly identify the low-frequency stimulus in the trials with more saccades (fixation: *t*(8) = 2.961, *p* = 0.018; pursuit: *t*(8) = 3.509, *p* = 0.008; paired *t*-tests). These trends were consistent across participants, with 8 out of 9 subjects showing improved performance during fixation and 7 subjects during pursuit. Thus, in both pursuit and fixation, retinal image motion caused by saccades and drifts strongly influences perception, differentially affecting visibility at low and high spatial frequencies.

## Discussion

Smooth pursuit eye movements aim to track moving objects, maintaining their image within the fovea for detailed inspection^4,5^. Consequently, retinal errors are often modeled as inputs for pursuit control^33,34^, leading to the assumption that target image motion on the retina during pursuit reflects tracking errors that the oculomotor system strives to eliminate. However, our findings suggest an alternative hypothesis: rather than minimizing retinal image motion, the pursuit system actively maintains controlled retinal “errors” for facilitation of visual processing. Crucially, two key observations support this view: 1) retinal image motion during pursuit substantially mimics that during fixation, and 2) these “errors” generate luminance transients that serve essential functional roles in vision during pursuit, analogous to their role in the saccade-fixation cycle. We discuss each in turn.

First, the spatiotemporal structure of retinal image motion during pursuit resembles that during fixation. If pursuit errors were merely tracking imperfections, such convergence would be unexpected. Yet, we observed striking similarities between pursuit- and fixation-induced retinal motion (Fig. **1**). The distribution of target image displacements on the retina is consistent across both tasks (Fig. **2**b), and retinal drift/saccade characteristics are remarkably alike (Fig. **2**c-e). These parallels persist not only in population averages but also as observer-specific traits across conditions (Figs. **3** and **4**). This alignment suggests a shared visual-oculomotor strategy and implementation involving controlled retinal motion.

Second, these retinal “errors” enhance visual processing. Akin to retinal motion during fixation^40,50–52^, pursuit-elicited retinal motion equalizes the strength of natural scenes within the peak frequency band of human spatial sensitivity (Fig. **2**f). This process reduces redundancies and enhances visual encoding efficiency^40,44,47,51,53^. Furthermore, pursuit-induced retinal motion shapes visual sensitivity. Consistent with reports of greater velocity variations during pursuit^54–56^, retinal drifts during pursuit are faster than during fixation (Fig. **5**). Consequently, pursuit increases luminance power in the human temporal sensitivity range at low spatial frequencies while attenuating power at high frequencies, leading to enhanced/reduced sensitivity at low/high frequencies (Fig. **6**). Critically, trials with greater luminance power during pursuit yield superior visual sensitivity (Fig. **7**). These perceptual modulations— absent under perfect retinal stabilization of the target image—indicate that pursuit actively maintains controlled retinal “errors” and exploits the resulting luminance transients to facilitate vision.

This hypothesis aligns with theories proposing temporal encoding of visual space. It has long been suggested that luminance modulations resulting from eye movements may contribute to spatial representations, particularly in high-acuity vision^19^. Recently these ideas have been expanded into the general proposal that visual encoding relies heavily on temporal processing^21^. Different types of eye movements exert distinct temporal modulations on static visual scenes and form spatiotemporal patterns with various frequency distributions^45,53^. Specifically, the oculomotor cycle of saccade/drift initiates coarse-to-fine processing^20,45^, with saccades enhancing low^50^ and drifts enhancing high^19,57,58^ spatial frequencies. Our study extends these space-time encoding theories to pursuit eye movements.

Our framework also reconciles with reported extra-retinal effects—specifically, visual suppression of low spatial frequency and enhancement of high spatial frequency during pursuit^30,31^. Schutz *et. al.* minimized retinal motion contributions using briefly flashed lines parallel to the pursuit direction and attributed the effects to extra-retinal facilitation. In contrast, our study minimized luminance transients unrelated to retinal motion using contrast-ramped stimuli embedded in the pursuit target. Since extra-retinal suppression preferably targets low spatial frequencies, like saccade and blink suppression^59^, it may mitigate perceived motion smear of untracked backgrounds^22,23^ but could interfere with target perception. Thus, the visual system may exploit the luminance modulations from target retinal motion to counteract extra-retinal influences.

This hypothesis implies that smooth pursuit simultaneously tracks and fixates on a moving target. “Tracking” maintains the target within the fovea, while “fixation” concurrently generates functional retinal motion of the target image. Thus, pursuit recruits an integrated process of tracking and processing of a moving target.

This view is supported in two ways. First, it resonates with the idea that pursuit is fixation of a moving target (or fixation is pursuit of a stationary target)^60^. For example, “fixation neurons” in the rostral superior colliculus encode foveal positional errors of visual targets during both fixation and pursuit^61^, echoing our observed retinal motion similarities. Additionally, the “gap effect” in fixation—the dependence of saccade latency on the interval between fixation target offset and saccadic target onset—persists during pursuit^62^, suggesting shared saccadic strategies. This aligns with our finding that saccades during pursuit resemble fixational saccades in characteristics and visual functions, implying that pursuit incorporates a fixational component applied to a moving target.

Second, pursuit’s “tracking” component aligns with its distinction from fixation, requiring disengagement^63^. Unlike fixation, where target perturbations minimally affect eye position, pursuit exhibits heightened sensitivity to such perturbations, with rapid eye velocity adjustments in response to target motion changes^64–66^. This enhanced sensitivity emerges at pursuit onset and vanishes when pursuit ceases^67^. In parallel, compared to steady fixation, visual tracking improves perceptual judgment of visual motion even with similar retinal stimulation^68^. Consistent with greater velocity variations^54–56^, we observed faster retinal drifts of the target image during pursuit than at fixation (Fig. **5**), potentially reflecting interference between tracking and fixation mechanisms.

Our study reveals a key open question: does the oculomotor system actively tune retinal motion during smooth pursuit—like during steady fixation—to optimize task-relevant signals? In fixation, ocular drifts adapt to the visual target’s spatial frequency^57^ and fine spatial details^69^, and fixational saccades optimize foveal positioning of visual images to counteract the nonuniform foveal sensitivity profile^70^. Indirect evidence hints at similar regulation during pursuit: predictive pursuit changes when identifying acuity targets^71^, and pursuit variability decreases when discriminating fine details within the moving stimulus^72^. Direct experimental validation remains essential in future work.

Finally, our findings highlight the need to consider eye movements in studies of visual impairment^73^. For example, Parkinson’s disease increases retinal displacement variance and saccade rates during both fixation and pursuit^74^, potentially impairing high-spatial-frequency signals and contributing to reduced acuity and contrast sensitivity to intermediate-to-high frequencies^75^. Similarly, autistic spectrum disorder involves greater retinal displacement variance and saccade amplitude during fixation^76,77^ and pursuit^77,78^, possibly contributing to facial processing difficulties^79^. Future research should investigate causal links between oculomotor deficits and visual impairments.

## Methods

### Subjects

Nine subjects (6 females and 3 males; average age 23 years) with emmetropic vision, as assessed with a standard Snellen eye-chart acuity test, participated in the study and received monetary compensation. All subjects, except one of the authors, were naive about the purposes of the study. The full study protocol was approved by the Research Subjects Review Board at the University of Rochester, and informed consent was obtained from every participant.

### Stimuli and Apparatus

Stimuli consisted of gray-scale Gabors (SD 0.8*^◦^*, spatial frequency either 1 or 10 cycles/degree, cpd) shown at the center of a disc, a circular region of uniform gray (diameter 4*^◦^*; luminance 12.8 cd*/*m^2^; Fig. **2**a). The stimuli were displayed over a slightly darker uniform background (luminance 12.0 cd*/*m^2^). The Gabors varied randomly across trials in their spatial frequency, orientation (±45*^◦^* relative to the vertical meridian), and phase (either 0*^◦^*, 90*^◦^*, 180*^◦^*, or 270*^◦^*). The stimuli either remained stationary on the monitor in the fixation condition or moved rightward at a uniform speed of 4 *^◦^/*s in the pursuit condition. The stimuli were masked by full-contrast plaids consisting of the 4 gratings at both frequencies (1 and 10 cpd) and orientations (±45*^◦^*). The mask was displayed over the disc (using the same Gaussian window of the stimulus, SD 0.8*^◦^*) in the late portion of the trial and over the entire screen at the end of the trial, as explained in the Procedure section.

Stimuli were presented on a calibrated LCD monitor (ASUS ROG SWIFT PG258Q) at a frame rate of 200 Hz and spatial resolution of 1920 × 1080 pixels. To faithfully render slight changes in contrasts on an eight-bit monitor, both the bit-stealing^80^ and the random dithering^81^ techniques were applied. Participants viewed the stimuli monocularly at a distance of 94 cm, with each pixel subtending approximately 1*^′^*. The observer’s head was immobilized via the use of a headrest and a custom-fabricated dental-imprint bite bar.

The movement of the right eye was continuously monitored by an in-house developed digital Dual Purkinje Image eye tracker (the dDPI^82^). This system yields sub-arcminute spatial resolution, as tested with artificial eyes, directly delivering eye position samples in digital format at a rate of 1,016 Hz. Examples of recorded eye trajectories are shown in Fig. **1**c and d.

### Experimental Procedure

Subjects were asked to either fixate on the target (in the fixation condition) or track it as it moved on the display (in the pursuit condition) and report the orientation of the Gabor. As shown in Fig. **2**a, in the pursuit trials, subjects initially fixated at a square dot (0.5*^◦^* wide, maximum contrast) which appeared at a visual angle of 5.4*^◦^* from the center of the monitor. After a random interval (400-600 ms) from the detection of fixation onset, the fixation marker disappeared and was replaced by the stimulus disc centered at the same location. Following a 500-ms delay, the disc started moving rightward at constant velocity. Then, 600 ms into the motion, the Gabor started to appear. Its contrast ramped up to an individually predetermined level over a period of 1.5 s before being masked by the plaid, which was displayed for 500 ms while the disc continued to move. A full-screen mask displayed for 1 s concluded the trial. Fixation trials consisted of the same sequence of stimuli presented at the center of the monitor and not moving.

The purpose of the contrast ramp was to minimize luminance modulations resulting from sources other than eye movements. The specific contrast value reached by the Gabor was predetermined for each subject at each spatial frequency. This was achieved via a preliminary calibration procedure using the PEST method^83^ to obtain ∼ 85% correct responses in the pursuit condition. Across subjects, the average ± SD of the contrast was 0.021 ± 0.004 with 1 cpd Gabors and 0.036 ± 0.013 with 10 cpd stimuli.

Data were collected in multiple experimental sessions each lasting approximately one hour. Each session consisted of the presentation of five blocks of trials. Within each block, 56 trials were randomly interleaved, with 14 repetitions for each of the 4 types of trials (1 or 10 cpd stimuli in fixation and pursuit). Before each block, the subject was comfortably seated with the head stabilized and the eye tracker was tuned for robust tracking of the Purkinje images.

To ensure accurate localization of the line of sight on the stimulus, rather than indirectly estimating gaze from oculomotor activity, as customary in the field, we directly estimated the center of gaze as perceived by the observer via a gaze-contingent procedure, as described in several previous publications^70,84,85^. In this procedure, following a standard 9-point calibration, subjects refine the estimated center of gaze that is displayed in real-time on the monitor by using a joypad. This approach has been shown to greatly improve the precision of gaze localization. Rests were taken by the subject in between blocks.

### Data Analysis

Only trials with continuous, uninterrupted eye-tracking were included for data analysis. Recorded eye traces were segmented into complementary periods of drifts and saccades in the fixation condition and periods of smooth pursuit and saccades in the pursuit condition. All the pursuit trials in which the subject failed to follow the stimulus in the period after the initial 300 ms of motion were excluded from data analysis. A pursuit failure was defined as a deviation between the eye and stimulus velocities larger than 60% for longer than 600 ms. As a result, the number of valid trials for an individual subject varied from 157 to 632 in every condition (pursuit or fixation at either high or low spatial frequency).

Due to the presence of pursuit eye movements, saccades cannot be detected directly based on the eye velocity. Instead, in both fixation and pursuit trials, we relied on the velocity of the stimulus on the retina, that is, the difference in velocity between the eye and the stimulus. In the fixation trials, since the stimulus did not move on the monitor, retinal motion was directly obtained by changing the sign of eye movements. Periods in which the retinal velocity was faster than 6 *^◦^/*s were labeled as saccades. The detecti results were confirmed and corrected as needed via visual inspection of all individual trials, as customary in the field^13,49^. We estimated the pursuit gain as the ratio *v*_e_*/v*_s_, where *v*_e_ and *v*_s_ respectively denote the velocities of the eye and stimulus during the pursuit intervals in between saccades.

In the pursuit trials, the segmentation of the reconstructed retinal signals yielded traces that closely resembled the drift-saccade alternation observed during fixation (Fig. **1**c and d). Therefore, for both fixation and pursuit trials, we refer to the complementary motion periods on the retina as retinal saccades and retinal drifts (or saccades and drifts, for brevity). Drift traces were smoothed using a 3rd-order Savitzty-Golay low-pass filter with a 41-ms window. The derivative of this filter was used to estimate drift velocity. Given our purpose of investigating the perceptual roles of retinal motion, we selectively focused on the period in which the Gabor stimulus was displayed (yellow shaded area in Fig. **2**a). In our analyses, since the retinal motion was virtually identical regardless of the spatial frequency of the stimulus, data collected with low (1 cpd) and high (10 cpd) spatial frequency Gabors were pooled together.

For both conditions, the probability density distribution of gaze position relative to the center of the stimulus was estimated over bins of 2.5*^′^*width for each individual (Fig. **3**a) and then averaged across subjects (Fig. **3**b). Distributions of instantaneous drift speed were estimated in bins of 2 *^′^*/s (Fig. **2**c). Diffusion constants, *D*, of drift motion were estimated by linear regression of the squared drift displacements against the time lag (also in Fig. **2**c; *D* = 1*/*4 of the regression slope). Direction distributions of drift segments and saccades in Figs. **2** and **4** were estimated with 10*^◦^* angular bins. Saccade amplitude and inter-saccadic interval distributions were estimated with 2*^′^* and 20-ms bins, respectively (Fig. **2**e).

To investigate the within-subject consistency of gaze positions relative to the stimulus across fixation and pursuit conditions, we employed linear regressions to assess the fixation-pursuit correlations in the statistical parameters of the gaze positions: horizontal (Fig. **3**d) and vertical means (Fig. **3**e), horizontal (Fig. **3**f) and vertical variances (Fig. **3**g), and horizontal-vertical covariance (Fig. **3**h).

To evaluate the resemblance in retinal drifts and saccades between fixation and pursuit at the individual level, we examined the correlations in individual characteristics between the two conditions. For each subject, we first obtained the circular mean directions of drift segments (Fig. **4**c), mean horizontal (Fig. **4**d) and vertical (Fig. **4**e) drift velocities, mean instantaneous drift speeds (Fig. **4**f), means of drift segments’ median curvatures (Fig. **4**g), circular mean saccades directions (Fig. **4**h), mean saccades amplitudes (Fig. **4**i), and mean inter-saccadic intervals (Fig. **4**j). Linear and circular^86^ regressions were then applied to evaluate the correlations in these mean characteristics between the two conditions. We also used paired *t*-tests across all individuals to quantitatively compare mean instantaneous drift speed (Fig. **5**a), mean of median curvatures (Fig. **5**b), diffusion constants (Fig. **5**c), mean saccade amplitudes and mean inter-saccadic intervals between conditions.

To study the link between oculomotor behavior and perceptual performance, we compared the proportions of correct responses in fixation and pursuit trials for both the low (1 cpd) and high (10 cpd) spatial frequency stimuli (Fig. **6**b). Paired *t*-tests and one-tailed Wilcoxon rank-sum tests were used to test statistical significance at the population and individual subject levels, respectively. To examine the visual consequences of saccade transients, we grouped trials according to whether their number of saccades was larger than the median and compared their performance (Fig. **7**d).

#### Similarity Score

To quantify the similarity between distributions of oculomotor parameters measured in the fixation and pursuit condition, we introduced a similarity score *S_JS_* based on the Jensen-Shannon distance *d_JS_*^87^. Given two distributions *p* and *q*,

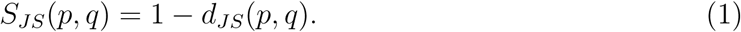

*d_JS_*(*p, q*) is a metric defined as the square root of the Jensen-Shannon divergence^88^:

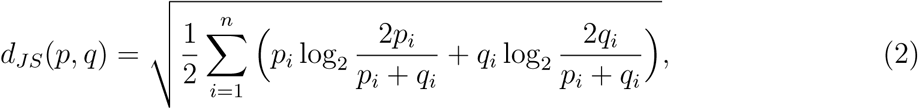

where *p_i_* and *q_i_* denote the discrete probabilities of the two respective distributions in the *i*th bin. The granularity of these probability distributions —i.e., the discrete intervals used to measure them — was taken to be the resolution with which the data were measured, as described in earlier paragraphs. Note that *d_JS_* ∈ [0, 1] (with 0 corresponding to identical distributions and 1 corresponding to non-overlapping distributions) and hence *S_JS_* ∈ [0, 1] (with 1 corresponding to identical distributions, and 0 corresponding to non-overlapping distributions). While the absolute size of *d_JS_*depends on this granularity, all of our comparisons were between distributions measured with equal granularity.

To determine the significance of within-subject vs. between-subject similarities, we applied two-sample *t*-tests for the within-subject similarity scores between fixation and pursuit against cross-subject scores between fixation and pursuit, between fixation and fixation, and between pursuit and pursuit (Fig. **3**b). The same analyses were also applied to the individual directional distributions of drift segments (Fig. **4**a) and saccades (Fig. **4**b).

We also compared how the gaze distribution of each individual varies from the population average for fixation and pursuit, by computing the distance (Eq. 2) of individual subjects’ data from the population average (excluding that subject) in each of the two conditions (Fig. **3**c) and then determining the correlation coefficients of these pairs of values (deviation for the fixation distribution and deviation for the pursuit distribution) across the subjects. We employed the same approach to compare, across the two conditions, the deviation of the gaze distribution from its 2D-Gaussian counterpart defined by the same parameters (Fig. **3**i).

#### Spectral Estimation

To examine the spatiotemporal signals delivered by fixation and pursuit, we reconstructed the luminance flow impinging onto the retina and estimated its power within the range of human temporal sensitivity. Given an image *I* as a function of space ***x***, the visual flow resulting from observing *I*(***x***) as it moves on the retina following a trajectory ***ξ***(*t*) can be expressed as

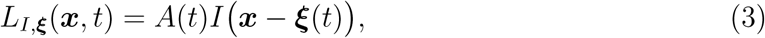

where *t* indicates time; *A*(*t*) represents the temporal modulations resulting from the time-varying contrast profile of the stimulus in our experiments (the 1.5-s ramp); and *I*(***x***) = sin (2*πk*_0_***x***) since we used a grating at spatial frequency *k*_0_ equal to 1 or 10 cycles/deg. In the fixation condition, the retinal image motion ***ξ***(*t*) is identical (opposite) to eye movements, whereas in the pursuit condition, ***ξ***(*t*) represents the difference between the subject’s eye movements and the stimulus’ trajectory.

The power spectrum of the visual input was estimated by the spatiotemporal factorization approach used in previous studies^40,89^:

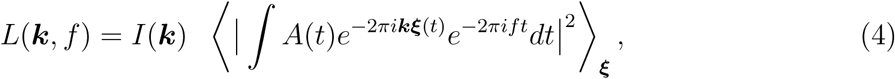

where ***k*** and *f* are the spatial and temporal frequencies and ⟨⟩***_ξ_*** indicates averaging across eye trajectories. The first term on the right side, *I*(***k***), is the spatial power spectrum of the stimulus, while the second term represents the temporal redistribution of power caused by both eye movements and the dynamics of stimulus presentation. This method enhances spectral resolution by assuming that the image on the retina and its motion are independent—a plausible assumption on average across the visual field.

To understand how this input signal may affect perception, we computed the integral of power within the temporal range of human sensitivity. That is, we first filtered the input signal by the known temporal sensitivity function of the human visual system^43^

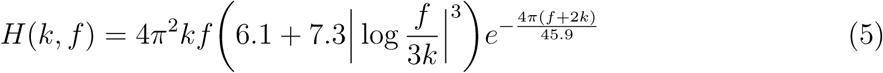

and then integrated across temporal frequencies:

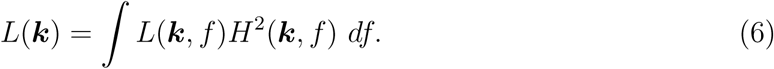

In Fig. **5**d and e, to examine how changes in retinal drift characteristics influence the redistribution of power on the retina, retinal drifts (the term ***ξ***(*t*) in Eq. 3) were modeled as Brownian-motion processes^40,41^ with varying diffusion constants (*D* = 20^57^, 30, 65, and 100 arcmin^2^/s). The term *A*(*t*) was here assumed to remain constant to represent stationary scenes. In the case of Brownian motion, the power redistribution caused by retinal drifts has a simple form^40^:

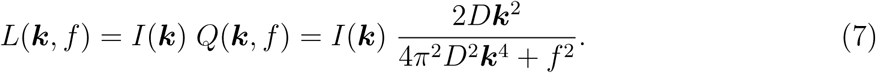

To compare the spatial-frequency dependence of the total power introduced by different drifts, we normalized the powers computed with Eqs. 5 and 7 by the value computed for the small diffusion constant (*D* = 30 arcmin^2^/s).

Similarly, to model how saccades influence the strength of the luminance signals impinging onto the retina, we simulated ocular drifts (*D* = 20 arcmin^2^/s) with and without saccadic intrusions (Fig. **7**a and b). Saccades were modeled using a sigmoid function in time (Eq. 8) with amplitude (*α*) and duration (*β*) randomly drawn from Gaussian distributions truncated at zero (*α* (arcmin) ∼ N(50.2, 27.0^2^); *β* (ms*^−^*^1^) ∼ N(0.291, 0.074^2^)) to match the experimentally observed distributions across observers. Each simulated saccade occurred in the middle of a drift trace, and its direction was randomly chosen from a uniform distribution of angles. Notably, confining the simulated saccades to horizontal directions had no impact on the results.

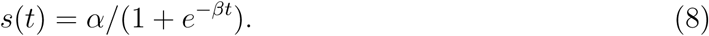

For every trial in the experiments of Fig. **2**a, we reconstructed the visual input experienced by the subject given the stimulus (*I* in Eq. 3), the individual contrast dynamics (*A*(*t*)), and the recorded eye trajectories (***ξ***(*t*)). Power spectra were estimated separately for each participant using all the available valid trials from the considered subject.

In Fig. **6**a, to examine the contribution of retinal drifts to perceptual performance, we conducted spectral analyses after removing saccades from the recorded eye traces. In this case, saccades were replaced with equal-length segments in which the eye remained immobile. Similarly, to assess the possible contributions from saccades, we computed power spectra after having assumed that the eye remained stationary during the recorded drift segments in between saccades (but used the actual saccade tracings, rather than the model of (Eq. 8) described above). The same approach was adapted in Fig. **2**f to investigate how retinal drifts and saccades reformat natural scenes (*I*(***k***) = 1*/****k***^2^); Here, *A*(*t*) = 1 was used to represent static scenes.

In Fig. **7**c, we examined how saccade occurrence influences low-frequency signal strength by dividing each subject’s fixation and pursuit trials into two groups (fewer vs. more saccades, based on the median) and computing the average power for each group using the recorded eye trajectories.

To emphasize how oculomotor activity influenced input power irrespective of the specific stimulus contrast used for each individual, the spectral measurements obtained from each participant in both Fig. **6**a and Fig. **7**c are shown after normalizing by the subject’s mean power. We used paired *t*-tests and one-tailed Wilcoxon rank-sum tests respectively to measure the significance of the power differences in the population data and each individual’s data.

## Acknowledgements

This study was supported by NIH grants R01 EY18363 (M.R.) and R01 EY07977 (J.D.V.). We thank Janis Intoy for helpful discussions on this work.

## Author Contributions

B.Y., J.D.V., and M.R. designed research and wrote the manuscript; B.Y. collected and analyzed data; and M.R. supervised the project.

